# The Auxin Response Factor ARF27 is required for maize root morphogenesis

**DOI:** 10.1101/2023.08.15.553439

**Authors:** Linkan Dash, Maxwell R. McReynolds, Melissa A. Draves, Rajdeep S. Khangura, Rebekah L. Muench, Jasper Khor, Jodi B. Callwood, Craig L. Cowling, Ludvin Mejia, Michelle G. Lang, Brian P. Dilkes, Justin W. Walley, Dior R. Kelley

## Abstract

Crop root systems are central to nutrition acquisition and water usage. Root hairs and lateral roots contribute to fine-scale patterning of root systems and can offer many advantages for improving root function without drastically impacting overall system architecture. Key genetic regulators underpinning root hair morphogenesis have been well characterized in the model plant Arabidopsis but are less understood in maize. Here, we identify a novel determinant of root hair morphogenesis and auxin responses in maize, AUXIN RESPONSE FACTOR27 (ARF27), using both reverse and quantitative genetic approaches. ARF27 is enriched in maize primary root tissues at both the transcript and protein level. Loss of *ARF27* leads to short primary roots and reduced root hair formation, while lateral root density is unaltered. In *arf27* roots, auxin-responsive gene expression is dysregulated, which is consistent with the predicted function of this transcription factor. Moreover, a genome wide association study (GWAS) to uncover genetic determinants of auxin-dependent root traits identified *ARF27* as a candidate gene. Furthermore, auxin hypersensitive maize genotypes exhibit altered crown root length and surface area in field-grown plants. A gene regulatory network (GRN) was reconstructed and an ARF27 subnetwork was integrated with DAP-seq and GWAS data to identify ARF27 target genes. The ARF27 GRN includes known maize root development genes, such as *ROOTLESS CONCERNING CROWN ROOTS (RTCS), ROOTHAIRLESS 3 (RTH3)* and *RTH6*. Altogether this work identifies a novel genetic driver of auxin-mediated root morphogenesis in maize that can inform agricultural strategies for improved crop performance.

## Introduction

Roots are a key organ for plant water and nutrient uptake. Root hairs are specialized epidermal cells critical for root function and have been implicated as critical structures for crop resilience^1–5^. The genetic networks underpinning root hair formation in Arabidopsis^6^ have been elucidated but are not as well understood in the model crop *Zea mays* (maize)^7^. Phytohormones, including auxin, brassinosteroids, cytokinin, and ethylene, act as influencers of Arabidopsis root hair formation via several transcription factor cascades, resulting in root hair initiation and initiation elongation^6^. In Arabidopsis, auxin can positively influence root hair morphogenesis via the DNA binding activity of conserved AUXIN RESPONSE FACTOR (ARF) transcription factors, ARF5, ARF7, and ARF19^8,9^. In maize, root hair formation has been linked to the activity of several *ROOTHAIRLESS (RTH)* genes^7^. *RTH1* encodes a SEC3 protein that is required for exocytotic vesicle fusion^10^. *RTH3* and *RTH6* have known roles in cellulose organization and synthesis, and encode a COBRA-like and CSLD5 proteins, respectively^11–13^. In addition, RTH5 is an NADPH oxidase required for maize root hair elongation^14^. Collectively these studies suggest that both cell wall and redox dynamics are important during maize trichoblast formation, but the transcriptional regulator(s) of this process are not yet known.

ARF transcription factors are conserved among land plants and have been well studied in Arabidopsis, *Marchantia polymorpha* and *Physcomitrium patens*^15^. The maize ARF family consists of 33 members, which display a high degree of overlap in transcriptional expression patterns^16–18^. Notably, protein abundance data for maize ARFs indicates that these transcription factors may exhibit some tissue specificity throughout development^19^. ARF family members fall into three clades designated A, B, and C^20^. DNA binding properties of maize ARFs across all three clades indicate a high degree of within clade overlap, suggesting that clade members may exhibit limited specificity^16^. In addition, Class A and Class B ARFs exhibited fairly distinct DNA binding landscapes^16^. A recent reconstruction of maize ARF gene regulatory networks in primary roots was in line with these findings^18^.

The unique roles of ARF genes in maize growth and development are emerging from molecular genetic studies. The evolutionarily conserved tasiR-ARF-ARF2/3 module is important for polarity establishment during leaf development^21^. During maize kernel development, ZmARF12 is required for repression of cell division, which is consistent with its designation as a Clade B ARF^22^. Members of all three clades have been implicated in maize root development: ARF4, ARF5, and ARF23^23–25^. While these studies demonstrate unique roles among ARF family members in maize, the functional characterization of the remaining 27 family members is needed to better understand these transcription factors.

Here, we report a novel role for maize *ARF27* in root hair development discovered using genetic approaches. Using a reverse genetic approach to discover ARF protein(s) with novel roles in maize root development, we identified ARF27 as a strong candidate based on its enriched protein abundance pattern in the elongation zone of primary roots^19^. Independently, a genome wide association study (GWAS) uncovered *ARF27* as a potential genetic determinant of auxin-dependent root growth. The maize *ARF27* gene is orthologous to Arabidopsis *AtARF7* and *AtARF19*^17^ and thus may represent an evolutionarily conserved regulatory module for root hair formation.

## Results

### Maize ARF27 is required for root hair formation

In order to determine ARF genes in maize that contribute to root morphogenesis, a reverse genetics screen was designed based on maize protein abundance atlas data^19^. Maize ARF proteins that exhibited enriched protein accumulation specifically in root tissues (primary root, elongation zone, meristematic zone, cortex, and/or seminal roots) included ARF3, ARF16, ARF19, ARF24, ARF27, ARF 30, ARF34^19^. For these ARF candidate genes, available transposon insertion alleles were ordered and characterized using established protocols, including validation of the transposons using locus-specific primers and backcrossing to the respective genotype to segregate away background insertions^26^. From this reverse genetic screen, ARF27 was identified as a strong candidate based on two key criteria. First, ARF27 exhibited enriched transcript expression in maize roots while protein was only detected in root tissues (Fig. S1A,B). Second, two UniformMu alleles of *ARF27* were heritable. The *arf27-1* allele (mu1078826) is located in the annotated promoter region of the gene while the *arf27-2* allele (mu1068742) is in the 5’ UTR (Fig. S1C; Table S1). Both alleles of *ARF27* are knock-downs and exhibit reduced expression of *ARF27* by RNA-seq and RT-qPCR analysis (Fig. S1D).

Phenotypic analysis of *arf27* loss of function alleles indicated that this transcription factor is required for several aspects of maize root morphogenesis. In the absence of *ARF27*, the root hairs are sparse and short in the elongation zone of *arf27-1* and *arf27-2* roots compared to wild-type W22 (Fig. 1A-C). In addition, exogenous treatment with 10 μM indole-3-acetic acid (IAA) leads to ectopic root hair formation within the meristematic zone in both alleles of *arf27* (Fig. 1D-I), suggesting an altered auxin response in this mutant. Loss of *ARF27* also leads to short primary roots (Fig. 1J), which are auxin insensitive (Fig. 1L). Both alleles of *arf27* have a normal lateral root density, irrespective of auxin treatment (Fig. 1K,L). Thus, *ARF27* is required for both primary root growth and root hair morphogenesis in maize.

**Fig. 1.**
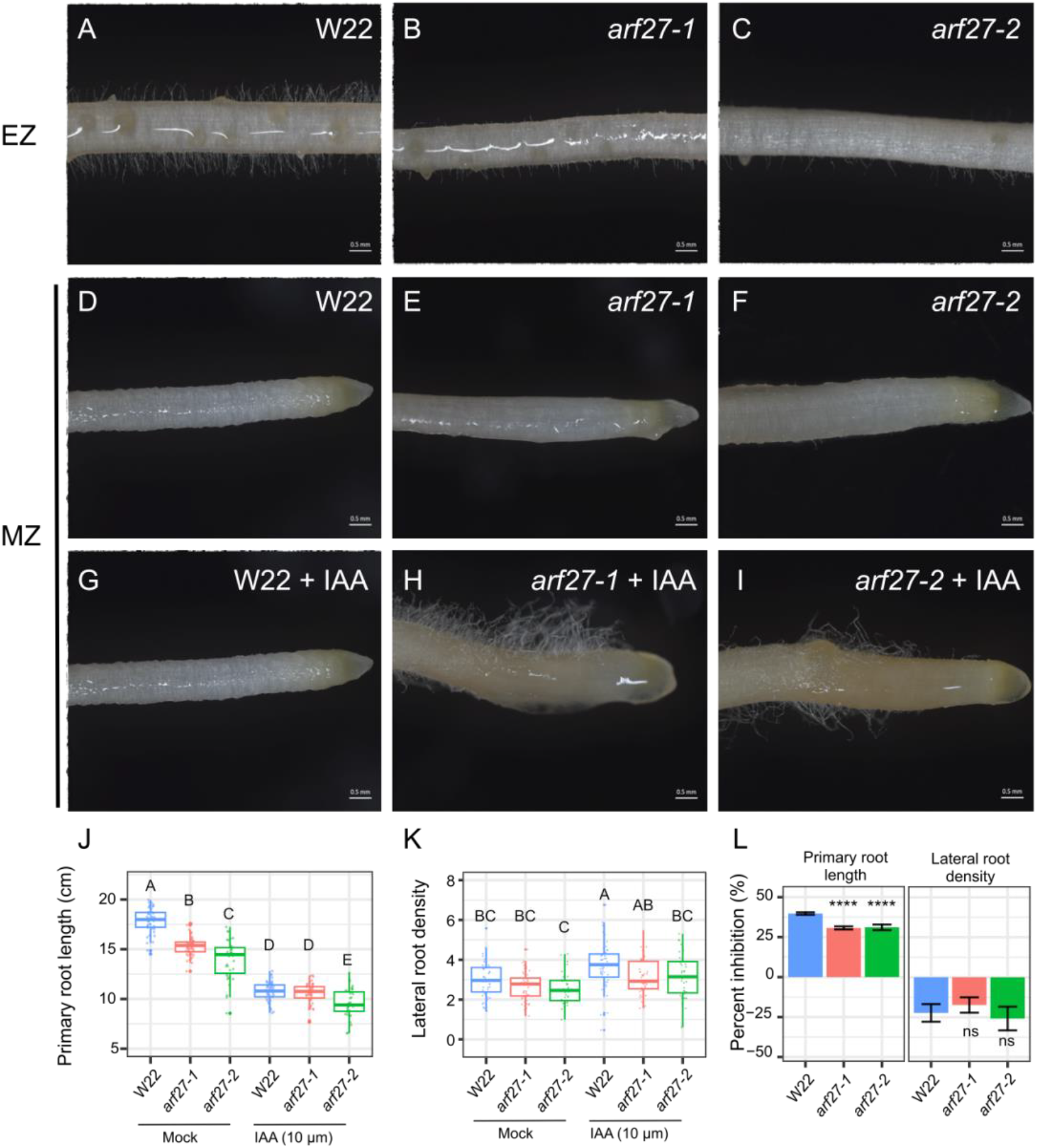
*ZmARF27* is required for maize root growth and development. Wild-type W22 root hairs form in the elongation zone (EZ) of primary roots (**A**) but not in the meristematic zone (MZ) (**D**). In *arf27-1* and *arf27-2* alleles, root hairs are sparse and short (**B, C**) and absent from the MZ (**E, F**). In the presence of exogenous auxin, ectopic root hairs form in the MZ of *arf27-1* and *arf27-2* (**H, I**) in contrast to W22 (**G**). Quantification of primary root length in 7-day-old seedlings in the presence and absence of 10 μM indole-3-acetic acid (IAA) (**J**). **(K)** Quantification of lateral root density (number of lateral roots per cm of the primary root) in 10-day-old seedlings in the presence and absence of 10 μM IAA. (**L**) Auxin-induced inhibition of root growth from (A) and auxin-induced increase in lateral root density from (B). The statistical test was one-way ANOVA, followed by Tukey’s post hoc test with a *P*<0.05. Letters indicate statistical significance of the differences observed in the data. Scale bars in A-I = 0.5 mm.

### ARF27 influences the auxin-dependent transcriptome in maize roots

ARF transcription factors are well known for influencing auxin-regulated gene expression in response to nuclear auxin signaling^15,16,27^. Transcriptional profiling of *arf27-1* and W22 roots was performed in the presence and absence of auxin treatment (10 μM IAA) to identify alterations in gene expression patterns in the mutant. Hundreds of auxin-responsive genes are misregulated in *arf27-1* roots compared to W22 (Fig. 2A,B; Table S2). In contrast, only 86 genes exhibit their normal auxin-responsive behavior in *arf27* roots (Fig. 2B). Roughly similar number of genes are up-and down-regulated in the absence of *ARF27* (Fig. 2A, B), which is consistent with another report on transcriptional changes associated with Class A ARFs from Arabidopsis^28^. Gene ontology (GO) enrichment analysis of ARF27-dependent transcripts indicates that auxin-activated signaling pathway genes are repressed in *arf27-1* compared to W22 (Fig. 2C). GO analysis of auxin-induced transcripts in *arf27-1* roots indicate that metabolism of trehalose, cytokinin, and gibberellin may be altered in response to auxin *arf27-1* roots (Fig. 2C). Auxin-repressed transcripts in *arf27-1* roots are enriched for cell wall associated GO terms (Fig. 2C). Collectively these data suggest that ARF27 is required for normal auxin-dependent transcriptional responses in maize roots.

**Fig. 2.**
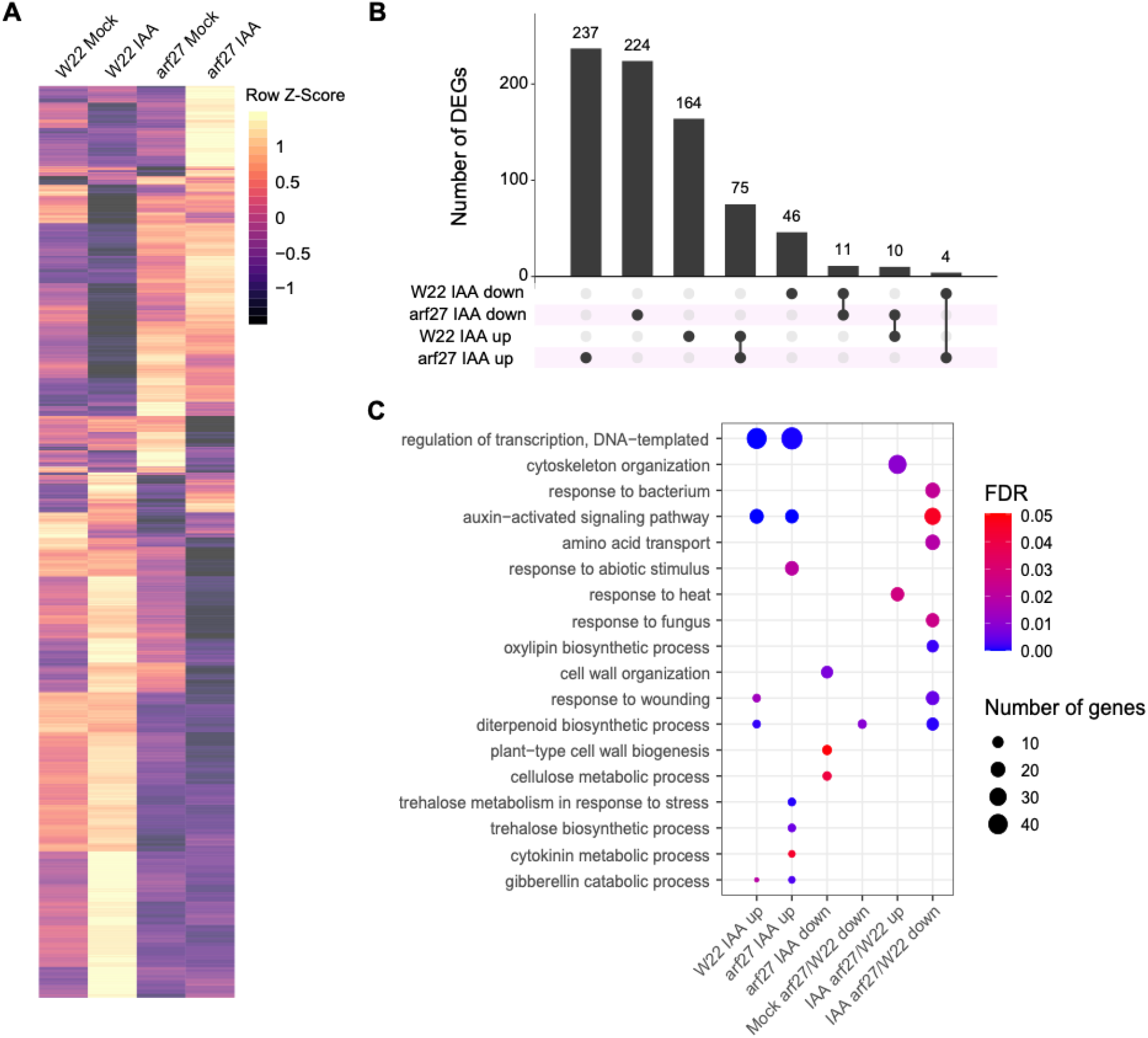
Auxin-responsive gene expression is altered in *arf27-1* roots. Hierarchical clustering of differentially expressed genes in W22 and *arf27-1* roots in the presence and absence of 1 hr 10 μM indole-3-acetic acid (IAA) treatment (**A**). Upset plot of DE genes indicates an overlap between transcripts across genotypes and treatments (**B**). Gene ontology enrichment of auxin-regulated DE genes in W22 and *arf27-1* (**C**).

### A genome wide association study to identify auxin-dependent root traits identifies ARF27

Genome wide association studies are a useful approach to identify candidate genes which contribute to phenotypes^29^. To complement the reverse genetic approach to identify key drivers of auxin-dependent root development in maize, we performed a genome wide association study (GWAS) using 617 genotypes from the Wisconsin Diversity Panel^30^. The selected 617 genotypes were divided into nine batches, with 72 genotypes per batch, based on a random complete block design. Every batch included B73 as a control reference genotype. Each genotype was grown for three days and then transferred to 0.5X Murashige-Skoog (MS) growth media (“mock”) or 0.5X MS supplemented with 10 μM indole-3-acetic acid (IAA) and grown for an additional four days. Three key traits were quantified on 7-day-old seedlings of each genotype (Table S3): primary root length without IAA (Fig. 3A), primary root length with IAA treatment (Fig. 3B), and root length ratio of auxin treated compared to mock (Fig. 3C). B73 seedlings exhibited the classical response of primary root growth inhibition following exogenous auxin treatment^18^ (Table S3). Similar to B73, most maize inbred genotypes also exhibited ∼74% inhibition following auxin treatment (Fig. 3C). Notably, several maize genotypes appear to be auxin insensitive (i.e., IAA/mock root length > 1.0) and a few genotypes were auxin hypersensitive (i.e., IAA/mock root length < 0.5) (Fig. 3C). Genome-wide association mapping identified numerous single nucleotide polymorphisms (SNPs) associated with each trait, and we identified 39 candidate genes with an association for one or more traits with a *P-*value £ 10^−6^ (Fig. 3D, Table S4). Notable hits among these candidate genes include *ARF27*, as well as two other *ARFs* (*ARF1* and *ARF16*) and *Zea mays SCARECROW* (*ZmSCR*) (Table S4). *Zm*SCR plays a key role in root morphogenesis and patterning^31–33^, demonstrating the validity of this GWAS approach. Furthermore, these data provide a suite of other top candidate genes that can be evaluated for their functional roles using reverse genetics approaches (Table S4).

**Fig. 3.**
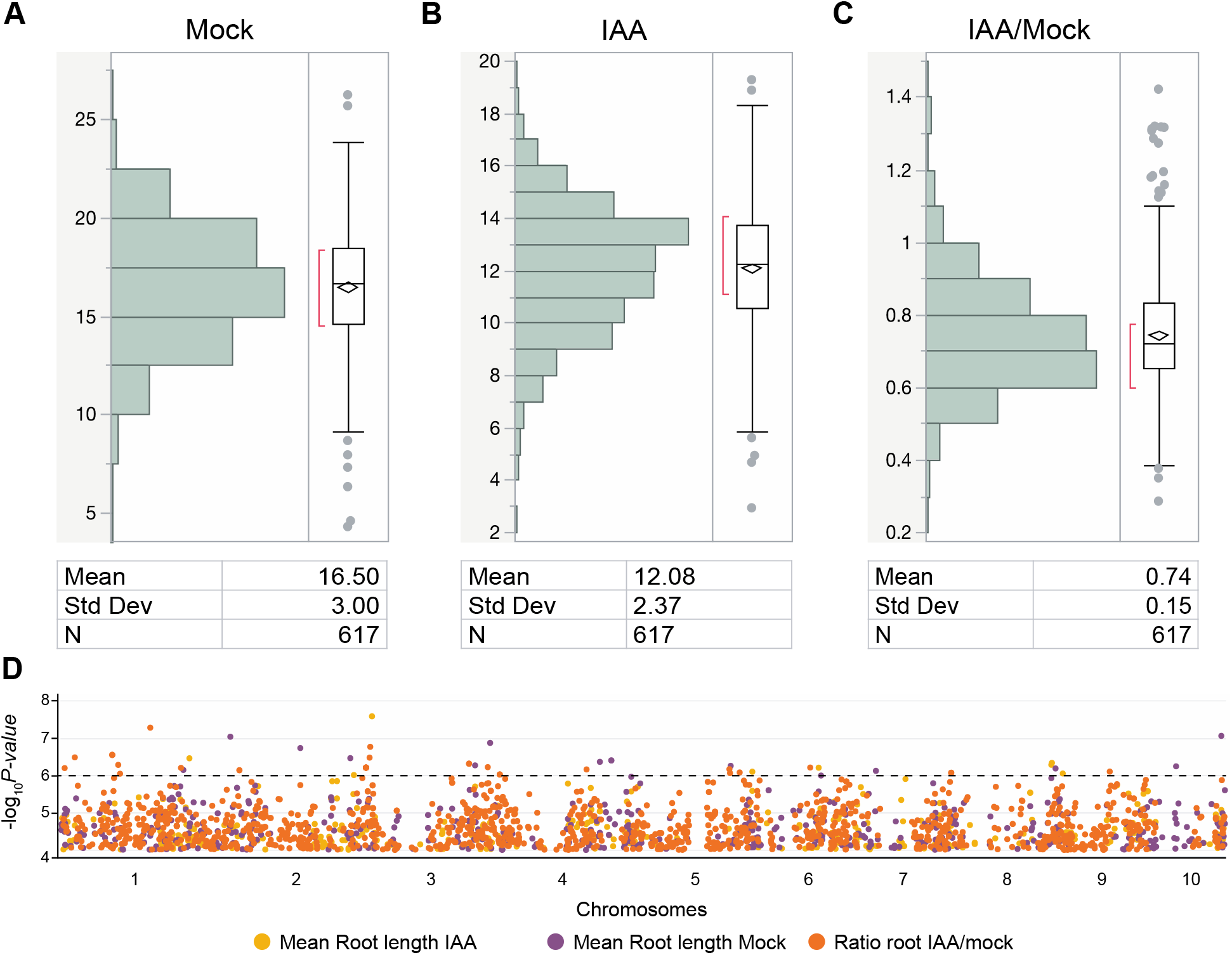
Genome wide association for maize root traits and auxin response. 7-day-old primary root length in 617 maize genotypes from the Wisconsin Diversity (WiDiv) Panel (**A**). 7-day-old primary root length in 617 WiDiv seedlings treated with 10 μM indole-3-acetic acid (IAA) for four days (**B**). Ratio of IAA-treated to mock-treated seedling root length in WiDiv lines (**C**). Genome-wide association mapping of >20 million single nucleotide polymorphisms (SNPs) for mean root length (purple dots), mean root length under IAA treatment (yellow dots), and ration of IAA/mock treated root length (orange dots) (**D**).

### Auxin hypersensitive maize genotypes exhibit altered crown root system architecture

Crown root systems are a key trait for maize physiology and several studies have quantified root system architecture using various approaches^34–36^. Among the WiDiv genotypes examined, three genotypes exhibited an hypersensitive auxin response compared to B73 (Fig. 3C). In order to determine if this phenotype was associated with crown root architecture traits, the top 20 outlier genotypes were grown in the field and characterized using the so-called “shovelomics” approach^37^. Three auxin hypersensitive genotypes (A401, DKIB02, and YONG 28), and two reference genotypes (B73 and W22) were phenotyped for numerous root traits using RhizoVision^38^ (Table S5). Notably, two of the auxin hypersensitive genotypes displayed reduced crown root systems compared to B73 (Fig. 4A-D). Specifically, both A401 and YONG 28 exhibited shorter crown root systems (Fig. 4E) and decreased total surface area compared to B73 (Fig. 4F). These findings suggest that genetic control of auxin response in maize root systems can influence mature crown root architecture under field conditions.

**Fig. 4.**
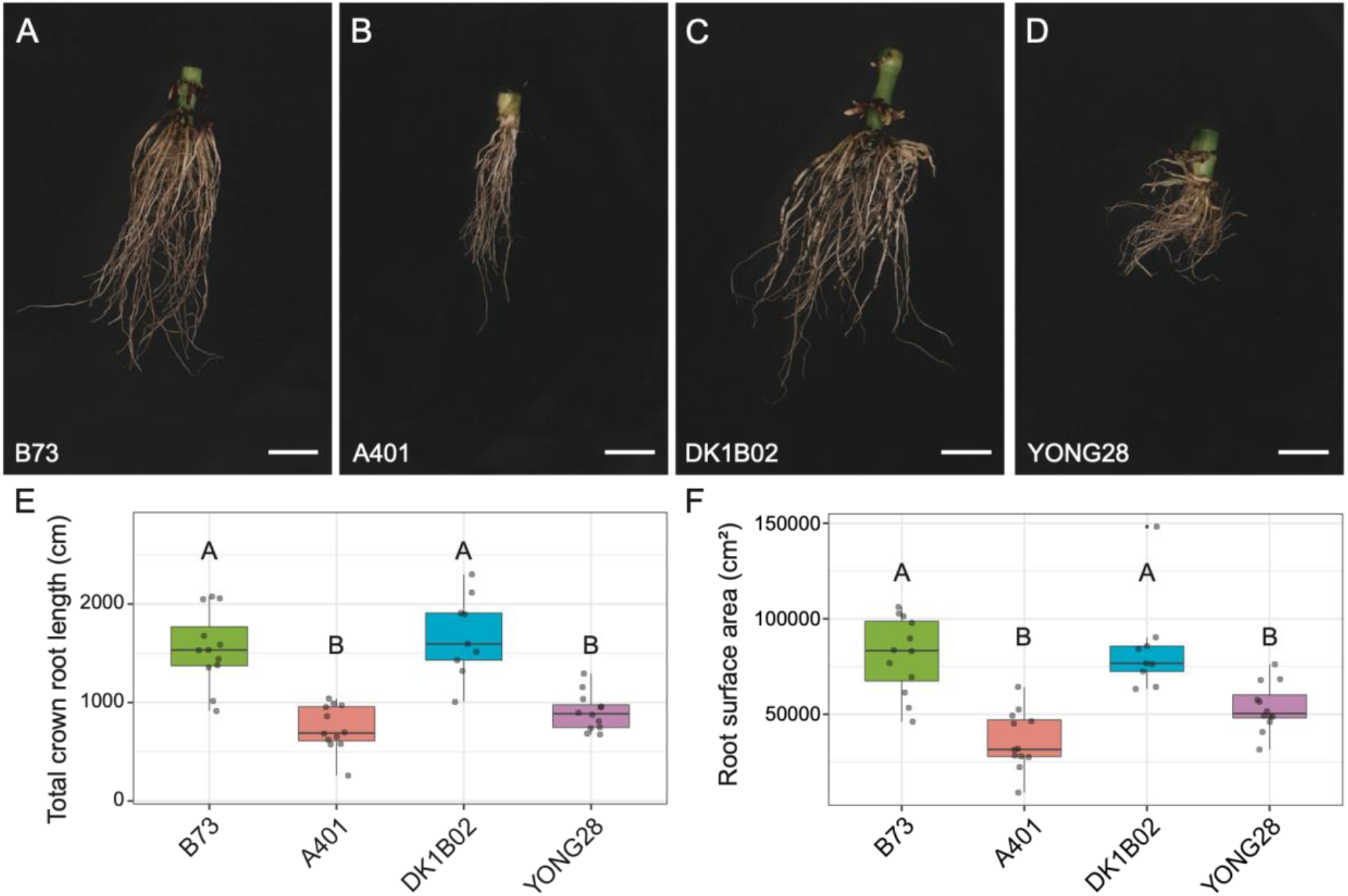
Crown root phenotypes of auxin hypersensitive genotypes. Representative images of field-grown crown root systems (**A**) B73 (**B**) A401 (**C**) DK1B02 (**D**) YONG 28 (**E**) Total crown root length calculated from 12 biological replicates per genotype. (**F**) Root surface area calculated from 12 biological replicates per genotype. In (E-F), statistical analysis was performed using a one-way ANOVA followed by Tukey’s posthoc analysis for multiple comparisons of means. The letters represent the statistical significance of the differences observed in the data. The images are digitally extracted for comparison, where all scale bars = 5.0 cm.

### Reconstruction of the maize ARF27 gene regulatory network

*ARF27* has been characterized as an auxin-responsive gene in maize, and it encodes for a putative activator transcription factor that binds a canonical Auxin Response Element (AuxRE) TGTC(GG) in maize ^1–3^. Putative target genes of maize ARF27 have been identified using DNA-affinity purification followed by sequencing (DAP-seq)^16^. We reconstructed a gene regulatory network (GRN) using Spatiotemporal Clustering and Inference of Omics Networks (SC-ION)^39^ based on the mock and IAA treated W22 and *arf27-1* transcript profiling data (Fig. 2; Table S2). We then extracted the first-neighbor targets of ARF27 comprised of 606 nodes and 648 edges (Fig. 5B; Table S6). Next, this subnetwork was overlaid with ARF27 DAP-seq targets^16^ and all root trait GWAS hits (Table S4) to examine the overlap across molecular data sets (Fig. 5A). In total, 46% of the SC-ION predicted ARF27 targets are also bound by ARF27 in DAP-seq assays (Fig. 5B, blue nodes). A subset of SC-ION predicted ARF27 targets were also among the ARF27 DAP-seq and GWAS hits (Fig. 5B, green nodes). Finally, a subset of the GRN predicted ARF27 targets are also GWAS hits (Fig. 5B, orange nodes). Among the ARF27 targets, several TFs are predicted to influence *ARF27* expression (Fig. 5B, dashed lines). Notable targets within the ARF27 network include well-characterized genetic regulators of maize root development: *ROOTLESS CONCERNING CROWN ROOTS* (*RTCS*)^40,41^, *RTCL-LIKE (RTCL1)*^41^, *ROOTHAIRLESS 3* (*RTH3*) and *RTH6*^11,13^ (Fig. 5B; Table S6). In addition, several Aux/IAA genes are putative ZmARF27 targets among early auxin-responsive genes bound by both Class A and Class B ARFs^16^.

**Fig. 5.**
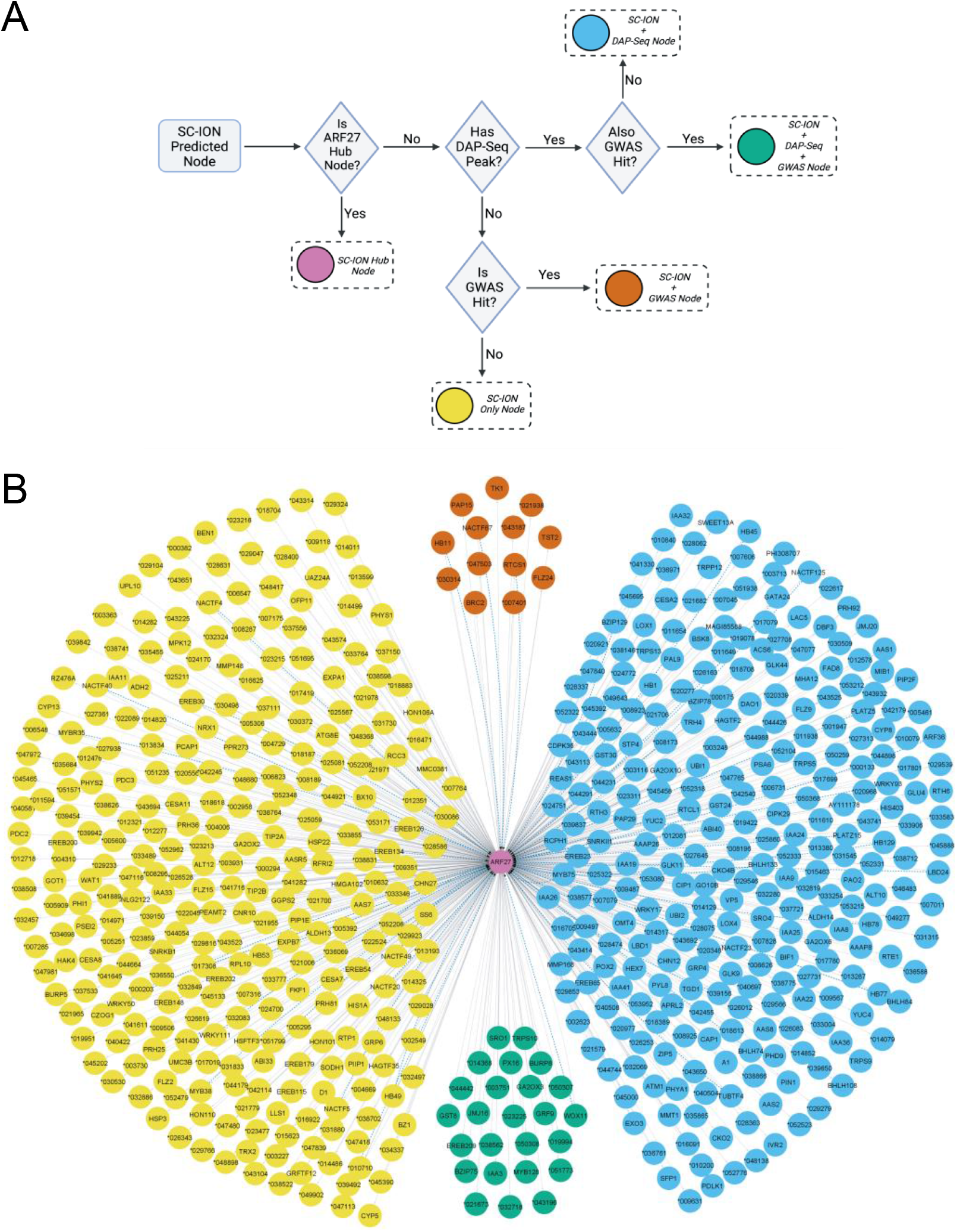
ZmARF27 gene regulatory network. Logic model for ZmARF27 network visualization. ZmARF27 is the central hub transcription factor (TF) indicated in pink. Targets predicted only by SC-ION are shown as yellow nodes. If a SC-ION predicted target is also present in the GWAS data, it is a vermillion-colored node. SC-ION targets which are also bound by DAP-seq data from Galli et al., 2018 are colored blue. Green nodes represent SC-ION targets which are also found in DAP-seq and GWAS data (**A**) ZmARF27 gene regulatory network with target nodes colored according to the logical model. Dashed lines represent TFs that are also predicted to regulate the expression of ZmARF27 (**B**).

## Discussion

Collectively, these genetic and molecular data support a novel role for ARF27 in regulating numerous aspects of maize root morphogenesis. Specifically, ARF27 is required for root hair formation and for primary root growth but it does not appear to play a role in lateral root formation. In the absence of *ARF27*, primary maize roots exhibit altered gene expression profiles and ectopic root hair formation. The Arabidopsis orthologs of maize ARF27, AtARF7 and AtARF19, are also positive regulators of root hair formation^6,42,43^. In addition, AtARF7 and AtARF19 have been linked to proper auxin-regulated gene expression, primary root growth, and lateral root formation^28,42–44^. This implies that these Class A ARFs may have evolutionarily conserved roles in root hair formation between monocots and eudicots but a divergence in lateral root function. However, it is possible that additional maize ARF genes are required for lateral root formation and a higher order mutant is required to uncover such roles. Notably, the establishment of root hair cell identity occurs via different mechanisms in maize and Arabidopsis^45^ suggesting divergent trichoblast formation cues between monocots and eudicots. Further characterization of *in vivo* bound and regulated genes for *Zm*ARF27, *At*ARF7, and *At*ARF19 will be required to determine the extent of overlap/conservation of their target genes.

The reduced expression of *ARF27* tracks with several desirable root traits and thus represents a potential target gene for genetic modification to improve root surface area. For example, maize mutants with reduced root architecture confer an advantage during water stress^46,47^ and thus ARF27-altered genotypes could be leveraged for improved drought tolerance or increased planting density. Despite the fact that root hairs increase surface area, the roles of root hairs in water uptake appears to be species specific and variable^48^. In maize, the root hair length is controlled by several different *RTH* genes, some of which may be direct in vivo targets of ARF27. Root hair length is a variable trait that can be influenced by phosphorous availability in Arabidopsis^49^, maize^50^, and barley^51^. Phosphate uptake rates vary among maize genotypes^52^ suggesting that this trait is under genetic control. Thus, future investigations into ARF27 as a breeding target for maize with improved phosphate uptake would be beneficial. In addition, the identification of several maize inbreds with diverse auxin responses (hyper-and insensitive genotypes) can be leveraged to better understand the impacts of auxinic herbicides on monocot crops^20,53^ and for informing genetic strategies to alter root system architecture^54,55^.

In summary, our results enhance our understanding of the transcriptional network operating in response to auxin in maize roots and how ARF27 modulates this response. Finally, the GWAS results describe the genetic architecture of the maize root response to auxin and identify numerous candidates for further study into their role in auxin-dependent root growth.

## Materials and Methods

### Plant materials

UniformMu seed stocks were obtained from the Maize Genetics Cooperation Stock Center and genotyped using gene-specific primers (Supplemental Table 4). Alleles were designated as *arf27-1* (mu1078826) and *arf27-2* (mu1068742). For phenotyping assays, maize seedlings were grown in rolled towel assays in a Percival growth chamber under long-day conditions (14 hours light, 10 hours dark) at 20ºC as previously described^56^. Before planting, the kernels were surfaced-sterilized with 50% bleach for 10 minutes and rinsed thrice with sterile water. Sterilized kernels were planted on seed germination paper (Anchor Paper company) pre-soaked with Captan fungicide. The rolled towels were placed vertically in 4L Nalgene beakers containing 600 mL of 0.5X Linsmaier and Skoog (LS) liquid media. Two days after planting, the rolls were opened and scored for germination; any ungerminated kernels were removed to synchronize seedling growth. For auxin response assays, three days after germination (DAG), the seedlings were placed in either 600 mL of 0.5X LS supplemented with 600 uL of 95% ethanol (“mock”) or 600 uL of 1 μM indole-3-acetic acid dissolved in 95% ethanol (“IAA”) for a final concentration of 10 μM IAA. Seedlings were allowed to grow for an additional 2-8 days and imaged at 5 DAG and 10 DAG. W22 and *arf27-1* seedlings were grown as described above for gene expression profiling experiments. On 5 DAG the seedlings were treated with 10 μM IAA or mock control for 1 hour and the primary roots were harvested from each seedling using a razor blade. Roots were pooled to generate replicates with 1 g of root tissue per genotype and treatment, then immediately flash-frozen in liquid nitrogen. Images of maize five-day-old primary roots were acquired with a Zeiss MacroZoom.

### Reverse Transcription-quantitative Polymerase Chain Reaction (RT-qPCR)

Five-day-old maize seedlings were grown using the rolled towel method. Root tissue from five seedlings was harvested and pooled together for each biological replicate, flash-frozen in liquid nitrogen, and ground to a homogenous fine powder using a mortar and pestle. Total RNA was extracted from ground root samples using TRIzol™ Reagent (Thermo Fisher Scientific, Catalog number 15596026), followed by purification using Zymo RNA MiniPrep kit (Fisher Scientific, Catalog number 50-444-628). The purified total RNA was used to synthesize cDNA using LunaScript® RT SuperMix Kit (New England Biolabs, Catalog number E3010). The synthesized cDNA was analyzed using qPCR using Luna® Universal qPCR Master Mix (NEB, catalog number M3003). qPCR data was analyzed as previously described^57^.

### QuantSeq and differential gene expression analysis

The differentially upregulated or downregulated genes were determined from QuantSeq data using PoissonSeq as previously described^18^. Differential expression was determined using a False Discovery Rate (FDR) <0.05. Upset plots were generated in R as previously described^18^.

### Gene ontology analysis

The differentially expressed genes were also tested for statistical over-representation of Gene Ontology terms for the biological processes in *Zea mays* using the Panther database as previously described^18^.

### Genome Wide Association Analyses

The Wisconsin Diversity (WiDiv) panel comprising 617 diverse inbred maize lines was used for the genome wide association study (GWAS). A random block design was used to group 72 genotypes per batch of growth assays, including a B73 check within each batch. For each genotype, 20 kernels were surfaced sterilized and planted on Captan-soaked seed germination paper using the rolled towel method^56^. Rolled towels were placed in beakers containing 0.5X MS media and kept in a controlled environment chamber at 20°C and 14h light/10h dark (long day) conditions. Seeds were scored for germination four days after planting, and ungerminated kernels were removed; this was designed “day 1”. Auxin treatments were performed as described^56^. Three days after germination (DAG), each genotype was split into either auxin media (10 μM indole-3-acetic acid in 0.5X MS) or control media (0.5X MS supplemented with an equal volume of 95% ethanol) and allowed to continue growing for an additional four days. Growth media was replaced at 5 DAG. Seedlings were imaged at 7 DAG using smartphone cameras. Primary root lengths were measured via hand annotation in Fiji with the segmented line tool for 5-10 individual seedlings per genotype and treatment (Table S3).

The mean trait values of the mock and IAA-treated maize seedling primary root length were used for GWAS. The marker genotypes of the 942 inbred lines from WiDiv consisting of 899,784 RNA-seq derived single nucleotide polymorphisms (SNPs) were obtained from Mazaheri et al., 2019^58^. These genotypes were imputed with the Beagle v5.1 software package^59^ using the HapMap3 SNPs^60^ as a reference panel. Imputations were performed using a window size of 30 Mb with a window overlap of 3.6 Mb, 10 burning, 15 iterations, and an effective population size of 1000. These fully imputed 55,242,281 SNP positions were filtered to retain SNPs with a minor allele frequency greater than 0.05 for the subset of 621 inbred lines from WiDiv used in this study. Filtering retained 22,507,869 SNPs that were used for GWAS analysis using the switchgrass R package^61^ modified for maize. The *P* value of 2.2 x 10^−9^ to identify statistically significant associations was determined for multiple-test correction using a Bonferroni-corrected threshold at α=0.05. To avoid false negative assessments of associations due to a conservative Bonferroni-corrected threshold, we analysed SNP associations at *P* value ≤ 10-^4^ for association analyses in this study. Using a custom Perl script we further annotated the lowest *P* value SNP in a 250 kb window with the nearest protein-coding gene (target), and two genes upstream and downstream of the target gene. This candidate list was used to prioritize genes for hypothesis testing.

### Crown Root Phenotyping (“Shovelomics”)

Twenty-two genotypes from the Wisconsin Diversity panel (including B73 and W22) were grown at the Iowa State University Ag Engineering & Agronomy Research farm during Summer 2022. Each genotype was grown in twin rows, with ten plants per row, in two separate nursery locations to minimize plot effects. The genotypes were planted according to a random complete block design to minimize row/genotype effects. At stage V6-V7, three plants per genotype and row were harvested to generate 12 biological replicates per genotype for image analysis. Individual plants were tagged with slip-n-lock vinyl labels for identification. All stem material above the third node was removed with Corona Forged steel 8” shears and discarded. Crown root systems were soaked in room temperature water for 10-30 minutes prior to washing with a commercial sink sprayer. Roots systems were air dried for six days prior to imaging. Images were acquired with a Canon E0S 80D DSLR camera with a 18-55 mm lens using a ring light and black vinyl photography background. Phenotype analysis was performed using RhizoVision^38^. Statistical significance was assessed using one-way ANOVA with a post-hoc Tukey’s test.

### Gene Regulatory Network reconstruction

The ZmARF27 GRN was reconstructed using SC-ION as previously described^18^ using all of the differentially expressed genes from Table S2. DAP-seq data was obtained from Galli et al., 2018^16^. The network was visualized in Cytoscape and color-coded using color-barrier-free annotations.

## Acknowledgments and funding sources

We dedicate this manuscript to Nick Lauter (1972 – 2021). Thank you to Addie Thompson (Michigan State University) for providing the initial Wisconsin Diversity panel source material (seed) in 2020. Thank you to Miriam Lopez (USDA ARS) and Marna Yandeau-Nelson for assistance with bulking the Wisconsin Diversity panel in 2020. This work was supported by United States Department of Agriculture (USDA) National Institute of Food and Agriculture (NIFA) Agriculture and Food Research Initiative (AFRI) grant 2020-67013-30914 to DRK and JWW; a USDA NIFA AFRI predoctoral grant 2021-67034-35188 to MRM; USDA NIFA AFRI postdoctoral grant 2022-^38^67012-36601 to RSK; Hatch Act and State of Iowa funds IOW03649 to DRK; and Hatch Act and State of Iowa funds IOW04108 to JWW.

## Author contributions

Conceptualization: MRM, LD, DRK, JWW, RSK, NL, BPD

Methodology: MRM, LD, MAD, MDL, RSK, BPD, JK, JBC

Investigation: MRM, MAD, MGL, RSK, CLC, JK, JBC, LM

Visualization: MRM, LD, RSK

Funding acquisition: DRK, JWW, RSK, BPD

Project administration: DRK, JWW Supervision: DRK, JWW, BPD

Writing – original draft: DRK, MRM, MAD, RSK, LD

Writing – review & editing: DRK, JWW

## Data and materials availability

All data, code, and genetic materials used in the analysis are available to any researcher for purposes of reproducing or extending the analysis. Transcriptome data is deposited at the Sequence Read Archive (SRA) as submission: SUB11800494 PRJNA858848. Seedling image data is available upon request. All other data are available in the main text or the supplementary materials.

## Supplementary Information

**Fig. S1.**
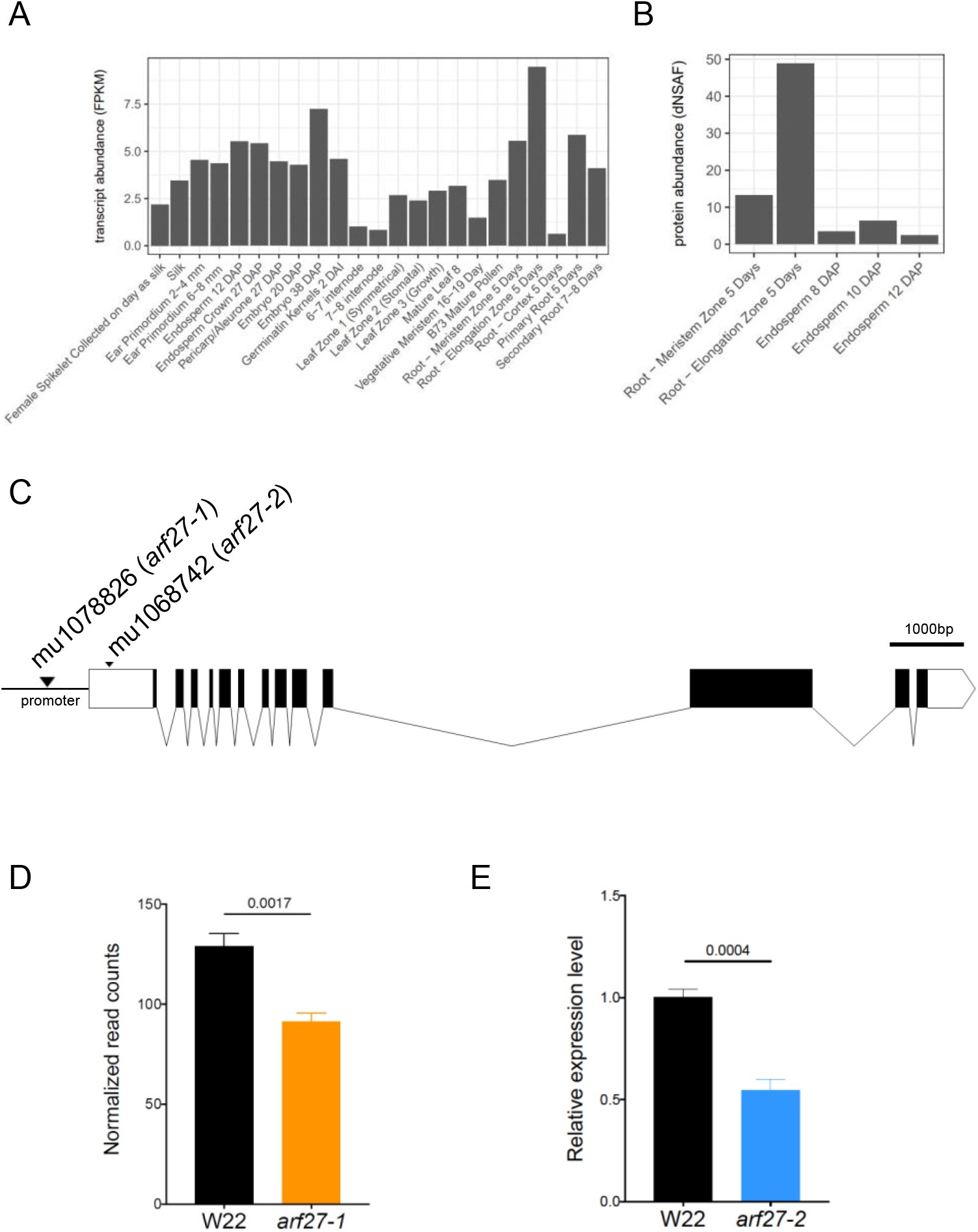
*ZmARF27* expression and allele characterization. (**A**) *ZmARF27* (Zm00001eb373970) is expressed throughout maize development with peak transcript abundance in the elongation zone of five-day-old roots. (**B**) ZmARF27 protein abundance is enriched in the elongation zone of five-day-old roots. (**C**) Gene model for *ZmARF27* with annotated UniformMu insertion alleles: *arf27-1* (mu1078826) and *arf27-2* (mu1068742). Untranslated regions (UTRs) are indicated in white boxes, exons are black boxes, and introns are lines. Scale bar = 1kb. (**D**) *ZmARF27* expression is reduced in *arf27-1*. (**E**) *ZmARF27* expression is reduced in *arf27-2. P* values are based on a t-test statistic.

**Table S1. Primers used for genotyping and RT-qPCR**. Oligonucleotide sequences for all primers used in this study.

**Table S2. Gene expression data in W22 and *arf27-1***. Normalized RNA-seq data for W22 and *arf27-1* roots treated with mock or 10 μM indole-3-acetic acid (IAA) for 1 hour, and differentially expressed (DE) genes between genotypes and/or treatments. GO enrichment analysis of DE genes.

**Table S3. Wisconsin Diversity panel seedling phenotype data**. Primary root length for seven-day-old maize inbreds grown the presence or absence of 10 μM indole-3-acetic acid (IAA or “auxin”) for four days.

**Table S4. Genome wide association study of maize primary root length with and without auxin treatment**.

**Table S5. Shovelomics phenotype data**. Root traits for 22 field-grown maize genotypes.

**Table S6. ZmARF27 gene regulatory network**. Predicted ZmARF27 targets reconstructed from an unsupervised approach overlaid with DAP-seq and GWAS hits.

## References

1. Ndoye, M. S., Burridge, J., Bhosale, R., Grondin, A. & Laplaze, L. Root traits for low input agroecosystems in Africa: Lessons from three case studies. Plant Cell & Environment 45, 637–649 (2022).

2. Rongsawat, T., Peltier, J.-B., Boyer, J.-C., Véry, A.-A. & Sentenac, H. Looking for Root Hairs to Overcome Poor Soils. Trends in Plant Science 26, 83–94 (2021).

3. Aslam, M. M., Karanja, J. K., Dodd, I. C., Waseem, M. & Weifeng, X. Rhizosheath: An adaptive root trait to improve plant tolerance to phosphorus and water deficits? Plant Cell & Environment 45, 2861–2874 (2022).

4. Hochholdinger, F. & Tuberosa, R. Genetic and genomic dissection of maize root development and architecture. Current Opinion in Plant Biology 12, 172–177 (2009).

5. Kohli, P. S., Maurya, K., Thakur, J. K., Bhosale, R. & Giri, J. Significance of root hairs in developing stress-resilient plants for sustainable crop production. Plant Cell & Environment 45, 677–694 (2022).

6. Vissenberg, K., Claeijs, N., Balcerowicz, D. & Schoenaers, S. Hormonal regulation of root hair growth and responses to the environment in Arabidopsis. Journal of Experimental Botany 71, 2412–2427 (2020).

7. Hochholdinger, F., Yu, P. & Marcon, C. Genetic Control of Root System Development in Maize. Trends in Plant Science 23, 79–88 (2018).

8. Mangano, S. et al. Molecular link between auxin and ROS-mediated polar growth. Proc. Natl. Acad. Sci. U.S.A. 114, 5289–5294 (2017).

9. Schoenaers, S. et al. The Auxin-Regulated CrRLK1L Kinase ERULUS Controls Cell Wall Composition during Root Hair Tip Growth. Current Biology 28, 722–732.e6 (2018).

10. Wen, T.-J., Hochholdinger, F., Sauer, M., Bruce, W. & Schnable, P. S. The roothairless1 Gene of Maize Encodes a Homolog of sec3, Which Is Involved in Polar Exocytosis. Plant Physiology 138, 1637–1643 (2005).

11. Li, L. et al. Characterization of maize roothairless6 which encodes a D-type cellulose synthase and controls the switch from bulge formation to tip growth. Sci Rep 6, 34395 (2016).

12. Penning, B. W. et al. Genetic Resources for Maize Cell Wall Biology. Plant Physiology 151, 1703–1728 (2009).

13. Hochholdinger, F. et al. The maize (Zea mays L.) roothairless3 gene encodes a putative GPI-anchored, monocot-specific, COBRA-like protein that significantly affects grain yield. Plant J 54, 888–898 (2008).

14. Nestler, J. et al. Roothairless5, which functions in maize (Zea mays L.) root hair initiation and elongation encodes a monocot-specific NADPH oxidase. Plant J 79, 729–740 (2014).

15. Li, Y., Han, S. & Qi, Y. Advances in structure and function of auxin response factor in plants. JIPB jipb.13392 (2022) doi:10.1111/jipb.13392.

16. Galli, M. et al. The DNA binding landscape of the maize AUXIN RESPONSE FACTOR family. Nat Commun 9, 4526 (2018).

17. Galli, M. et al. Auxin signaling modules regulate maize inflorescence architecture. Proc Natl Acad Sci U S A 112, 13372–13377 (2015).

18. McReynolds, M. R. et al. Temporal and spatial auxin responsive networks in maize primary roots. http://biorxiv.org/lookup/doi/10.1101/2022.02.01.478706 (2022) xdoi:10.1101/2022.02.01.478706.

19. Walley, J. W. et al. Integration of omic networks in a developmental atlas of maize. Science 353, 814–818 (2016).

20. Matthes, M. S. et al. Auxin EvoDevo: Conservation and Diversification of Genes Regulating Auxin Biosynthesis, Transport, and Signaling. Molecular Plant 12, 298–320 (2019).

21. Dotto, M. C. et al. Genome-Wide Analysis of leafbladeless1-Regulated and Phased Small RNAs Underscores the Importance of the TAS3 ta-siRNA Pathway to Maize Development. PLoS Genet 10, e1004826 (2014).

22. Wang, Y. et al. Transcriptomic Variations and Network Hubs Controlling Seed Size and Weight During Maize Seed Development. Front. Plant Sci. 13, 828923 (2022).

23. Li, J. et al. Maize Transcription Factor ZmARF4 Confers Phosphorus Tolerance by Promoting Root Morphological Development. IJMS 23, 2361 (2022).

24. Yang, F., Shi, Y., Zhao, M., Cheng, B. & Li, X. ZmIAA5 regulates maize root growth and development by interacting with ZmARF5 under the specific binding of ZmTCP15/16/17. PeerJ 10, e13710 (2022).

25. Liang, T. et al. GWAS across multiple environments and WGCNA suggest the involvement of ZmARF23 in embryonic callus induction from immature maize embryos. Theor Appl Genet 136, 93 (2023).

26. Settles, A. M. et al. Sequence-indexed mutations in maize using the UniformMu transposon-tagging population. BMC Genomics 8, 116 (2007).

27. Mutte, S. K. et al. Origin and evolution of the nuclear auxin response system. eLife 7, e33399 (2018).

28. Powers, S. K. et al. Nucleo-cytoplasmic Partitioning of ARF Proteins Controls Auxin Responses in Arabidopsis thaliana. Molecular Cell 76, 177–190.e5 (2019).

29. Tibbs Cortes, L., Zhang, Z. & Yu, J. Status and prospects of genome-wide association studies in plants. Plant Genome 14, (2021).

30. Hansey, C. N., Johnson, J. M., Sekhon, R. S., Kaeppler, S. M. & Leon, N. Genetic Diversity of a Maize Association Population with Restricted Phenology. Crop Sci. 51, 704–715 (2011).

31. Lim, J. et al. Conservation and Diversification of SCARECROW in Maize. Plant Mol Biol 59, 619–630 (2005).

32. Lim, J. et al. Molecular Analysis of the SCARECROW Gene in Maize Reveals a Common Basis for Radial Patterning in Diverse Meristems. Plant Cell 12, 1307–1318 (2000).

33. Hughes, T. E. & Langdale, J. A. SCARECROW is deployed in distinct contexts during rice and maize leaf development. Development 149, dev200410 (2022).

34. Ren, W. et al. Genome-wide dissection of changes in maize root system architecture during modern breeding. Nat. Plants 8, 1408–1422 (2022).

35. Liu, S., Barrow, C. S., Hanlon, M., Lynch, J. P. & Bucksch, A. DIRT/3D: 3D root phenotyping for field-grown maize (Zea mays). Plant Physiology 187, 739–757 (2021).

36. Zheng, Z. et al. Shared Genetic Control of Root System Architecture between Zea mays and Sorghum bicolor. Plant Physiol. 182, 977–991 (2020).

37. Trachsel, S., Kaeppler, S. M., Brown, K. M. & Lynch, J. P. Shovelomics: high throughput phenotyping of maize (Zea mays L.) root architecture in the field. Plant Soil 341, 75–87 (2011).

38. Seethepalli, A. et al. RhizoVision Crown: An Integrated Hardware and Software Platform for Root Crown Phenotyping. Plant Phenomics 2020, 1–15 (2020).

39. Clark, N. M. et al. Integrated omics networks reveal the temporal signaling events of brassinosteroid response in Arabidopsis. Nat Commun 12, 5858 (2021).

40. Muthreich, N. et al. Regulation of the maize (Zea mays L.) embryo proteome by RTCS which controls seminal root initiation. European Journal of Cell Biology 89, 242–249 (2010).

41. Taramino, G. et al. The maize (Zea mays L.) RTCS gene encodes a LOB domain protein that is a key regulator of embryonic seminal and post-embryonic shoot-borne root initiation: Map-based cloning of the maize RTCS gene. The Plant Journal 50, 649–659 (2007).

42. Bhosale, R. et al. A mechanistic framework for auxin dependent Arabidopsis root hair elongation to low external phosphate. Nat Commun 9, 1409 (2018).

43. Okushima, Y. et al. Functional Genomic Analysis of the AUXIN RESPONSE FACTOR Gene Family Members in Arabidopsis thaliana : Unique and Overlapping Functions of ARF7 and ARF19. The Plant Cell 17, 444–463 (2005).

44. Huang, K.-L. et al. The ARF7 and ARF19 Transcription Factors Positively Regulate PHOSPHATE STARVATION RESPONSE1 in Arabidopsis Roots. Plant Physiol 178, 413–427 (2018).

45. Marzec, M., Melzer, M. & Szarejko, I. Root Hair Development in the Grasses: What We Already Know and What We Still Need to Know. Plant Physiol. 168, 407–414 (2015).

46. Sebastian, J. et al. Grasses suppress shoot-borne roots to conserve water during drought. Proc. Natl. Acad. Sci. U.S.A. 113, 8861–8866 (2016).

47. Gao, Y. & Lynch, J. P. Reduced crown root number improves water acquisition under water deficit stress in maize (Zea mays L.). EXBOTJ 67, 4545–4557 (2016).

48. Cai, G. & Ahmed, M. A. The role of root hairs in water uptake: recent advances and future perspectives. Journal of Experimental Botany 73, 3330–3338 (2022).

49. Bates, T. R. & Lynch, J. P. Stimulation of root hair elongation in Arabidopsis thaliana by low phosphorus availability. Plant Cell Environ 19, 529–538 (1996).

50. Zhu, J., Kaeppler, S. M. & Lynch, J. P. Mapping of QTL controlling root hair length in maize (Zea mays L.) under phosphorus deficiency. Plant Soil 270, 299–310 (2005).

51. Haling, R. E. et al. Root hairs improve root penetration, root–soil contact, and phosphorus acquisition in soils of different strength. Journal of Experimental Botany 64, 3711–3721 (2013).

52. Griffiths, M. et al. A multiple ion-uptake phenotyping platform reveals shared mechanisms affecting nutrient uptake by roots. Plant Physiology 185, 781–795 (2021).

53. Johnson, N., Zhang, G., Soble, A., Johnson, S. & Baucom, R. S. The consequences of synthetic auxin herbicide on plant–herbivore interactions. Trends in Plant Science S136013852300050X (2023) doi:10.1016/j.tplants.2023.02.003.

54. Zhan, A., Schneider, H. & Lynch, J. P. Reduced Lateral Root Branching Density Improves Drought Tolerance in Maize. Plant Physiol 168, 1603–1615 (2015).

55. Zhan, A. & Lynch, J. P. Reduced frequency of lateral root branching improves N capture from low-N soils in maize. J Exp Bot 66, 2055–2065 (2015).

56. Draves, M. A., Muench, R. L., Lang, M. G. & Kelley, D. R. Maize Seedling Growth and Hormone Response Assays Using the Rolled Towel Method. Current Protocols 2, (2022).

57. Dash, L. et al. slim shady is a novel allele of PHYTOCHROME B present in the T-DNA line SALK_015201. http://biorxiv.org/lookup/doi/10.1101/2021.02.12.430994 (2021) xdoi:10.1101/2021.02.12.430994.

58. Mazaheri, M. et al. Genome-wide association analysis of stalk biomass and anatomical traits in maize. BMC Plant Biol 19, 45 (2019).

59. Browning, B. L., Zhou, Y. & Browning, S. R. A One-Penny Imputed Genome from Next-Generation Reference Panels. The American Journal of Human Genetics 103, 338–348 (2018).

60. Bukowski, R. et al. Construction of the third-generation Zea mays haplotype map. GigaScience 7, (2018).

61. Lovell, J. T. et al. Genomic mechanisms of climate adaptation in polyploid bioenergy switchgrass. Nature 590, 438–444 (2021).

